## Supplementary Info for "Phase-separating pyrenoid proteins form complexes in the dilute phase"

[8] Present address: Laboratory of Chemical Physics, National Institute of Diabetes and Digestive and Kidney Diseases, National Institutes of Health, Bethesda, Maryland 20892, USA

**Supplementary Note 1**

Although we managed to minimize EPYC1 sticking to the surface by using the PEI-PEG coating, we were not able to completely eliminate Rubisco sticking in the experiments described in Fig. 2. We thus expect the real concentration of Rubisco in solution to be lower than the nominal concentration. We added horizontal error bars in Fig. 2e to reflect this uncertainly in Rubisco concentrations. Specifically, we assume that Rubisco adsorption saturates at about 10 nM.

**Supplementary Note 2**

To estimate the $K_D$, we used the quadratic binding equation below to fit the data in Fig. 2d. This is because in our binding assay, Rubisco's concentration is not significantly higher than EPYC1's, so we cannot assume that only a small fraction of Rubisco bound to EPYC1[1]. The fraction of Rubisco bound EPYC1 can be expressed as

$$\frac{[\mathrm{AB}]}{A_0} = \frac{(A_0+B_0+K_\mathrm{d}) - \sqrt{(A_0+B_0+K_\mathrm{d})^2 - 4A_0B_0}}{2A_0}$$

In our case:

AB: Rubisco-EPYC1 complex

$A_0$: Rubisco concentration

$B_0$: EPYC1-GFP concentration

Then, diffusion rates of EPYC1-GFP measured by FCS should be weighted average of unbound and bound EPYC1-GFP, as $D_{\mathrm{EPYC1}}$ and $D_{\mathrm{complex}}$

$$D = \frac{[\mathrm{AB}]}{A_0} \cdot D_{\mathrm{complex}} + \left(1 - \frac{[\mathrm{AB}]}{A_0}\right) D_{\mathrm{EPYC1}}$$

$$= D_{\mathrm{complex}} \cdot \frac{(A_0+B_0+K_\mathrm{d}) - \sqrt{(A_0+B_0+K_\mathrm{d})^2 - 4A_0B_0}}{2A_0}$$

$$+D_{\mathrm{EPYC1}} \cdot (1 - \frac{(A_0+B_0+K_\mathrm{d}) - \sqrt{(A_0+B_0+K_\mathrm{d})^2 - 4A_0B_0}}{2A_0})$$

We used this equation to fit the data in Fig. 2d to extract $K_{\mathrm{d}}$ (unit: nM). We fixed $D_{\mathrm{EPYC1}} = 62\,\mu\mathrm{m}^2/\mathrm{s}$ and $D_{\mathrm{complex}} = 41\,\mu\mathrm{m}^2/\mathrm{s}$ to obtain the estimate of $K_{\mathrm{d}} = 29 \pm 12$ nM (error bars represent 68% confidence interval):

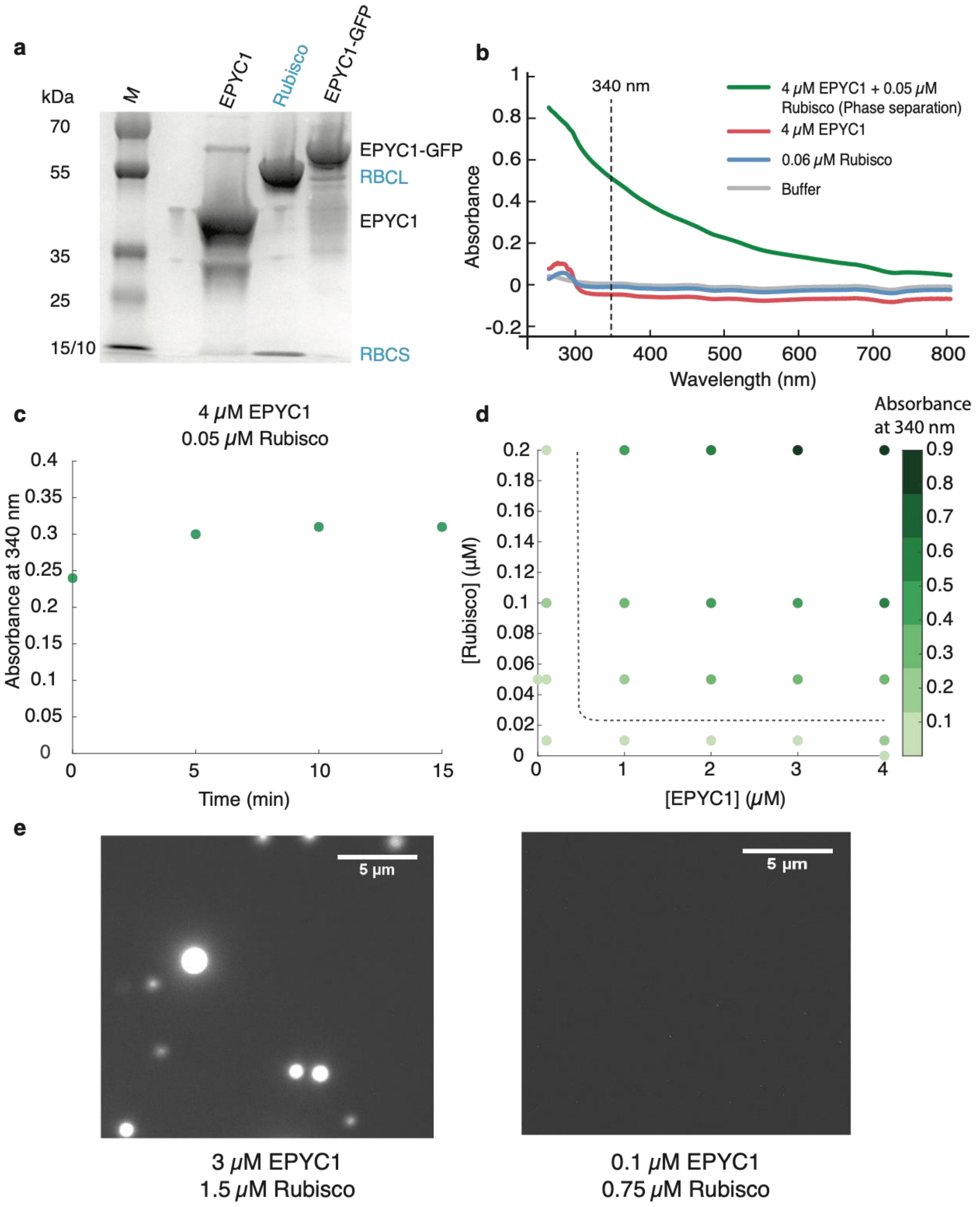


**Supplementary Figure 1 | Use of a turbidity assay and fluorescence imaging to determine the phase diagram of EPYC1 and Rubisco. a** Purified proteins on an SDS-PAGE gel. RBCL:

Rubisco large subunit. RBCS: Rubisco small subunit. M: marker. **b** UV/Vis extinction spectra of different sample solutions (the phase separated sample was measured 10 minutes after mixing). **c** The absorbance at 340 nm of a phase-separating solution consisting of 4 $\mu$M EPYC1 and 0.05 $\mu$M Rubisco as a function of time after mixing. **d** Absorption assay at 340 nm of mixed EPYC1-Rubisco solutions at concentrations shown on the *x* and *y* axes. **e** Representative fluorescence images of phase separation (left) and no phase separation (right). 20 nM EPYC1-GFP was added to both solutions.

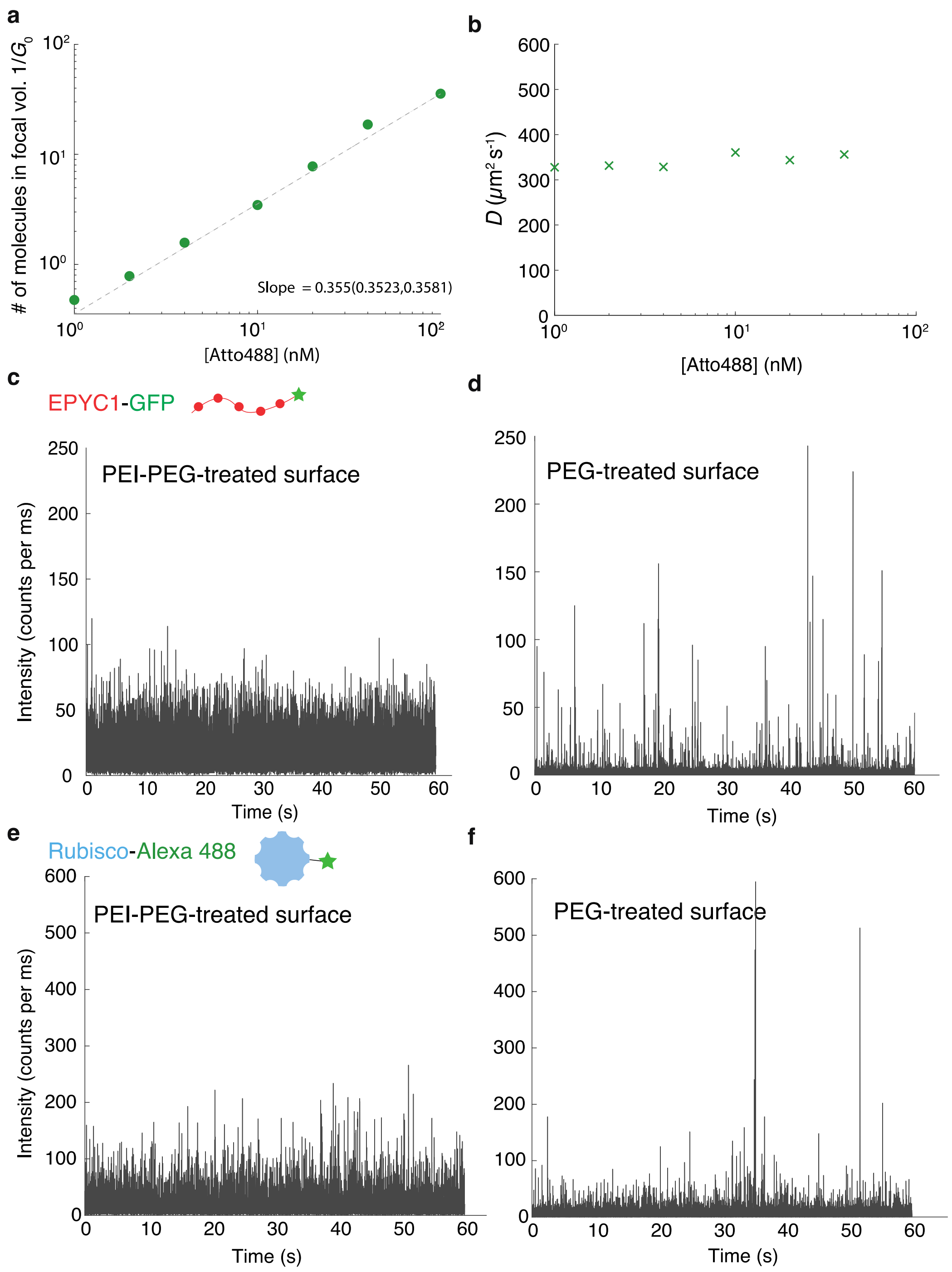


**Supplementary Figure 2 | FCS experiments were performed using a well-calibrated setup and an optimized surface. a** The number of Atto488 free dye molecules in the FCS focal volume were measured as a function of Atto488 concentrations. The focal volume was then calculated based on the slope of the fitted curve. **b** Atto488 diffusion rates were measured as a function of

Atto488 concentrations. **c-d** FCS raw trace of 10 nM EPYC1-GFP on PEI-PEG-treated surface (**c**) and PEG-treated surface (**d**)**. e-f** FCS raw trace of 10 nM Rubisco-Alexa 488 on PEI-PEG-treated surface (**e**) and PEG-treated surface (**f**). In panels **d** and **f**, the big spikes are protein aggregates, which were absent for PEI-PEG treated surfaces (panels **c** and **e**)

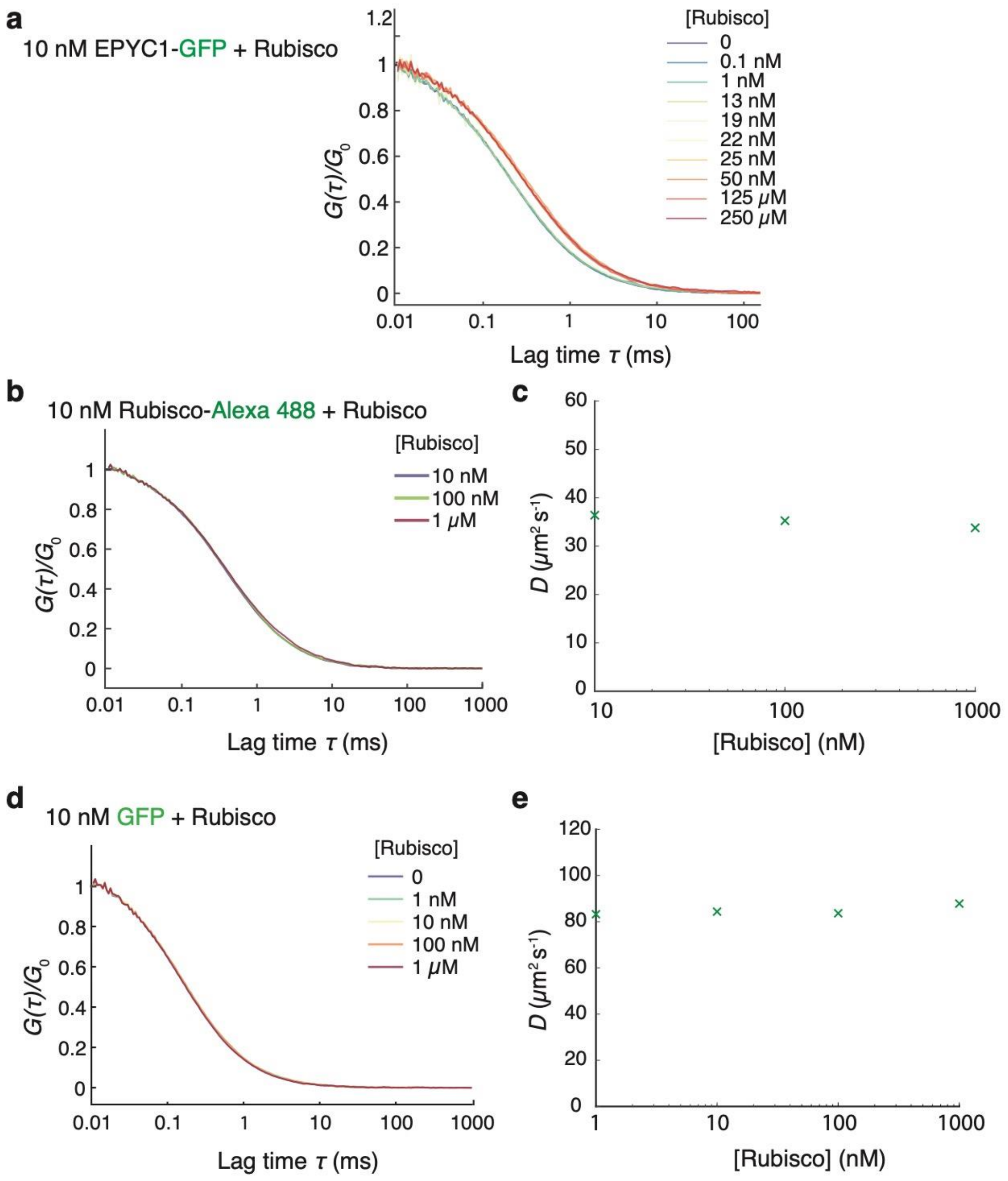


**Supplementary Figure 3 | Neither interactions between GFP and Rubisco nor Rubisco self-interactions were observed in the FCS experiments. a** Full autocorrelation curves of EPYC1-GFP fluorescence at different Rubisco concentrations. In **a**, **b**, **d**, each autocorrelation curve is normalized by its fitted value at $t$ = 0 (Methods). **b** Autocorrelation curves of Rubisco labelled with Alexa Fluor 488 at different unlabeled Rubisco concentrations. **c** Diffusion rates of Rubisco-

Alexa 488 inferred from FCS data in **b** as a function of total Rubisco concentration. **d** Autocorrelation curves of GFP at different unlabeled Rubisco concentrations. **e** Diffusion coefficients of GFP inferred from FCS data in **d** as a function of Rubisco concentration.

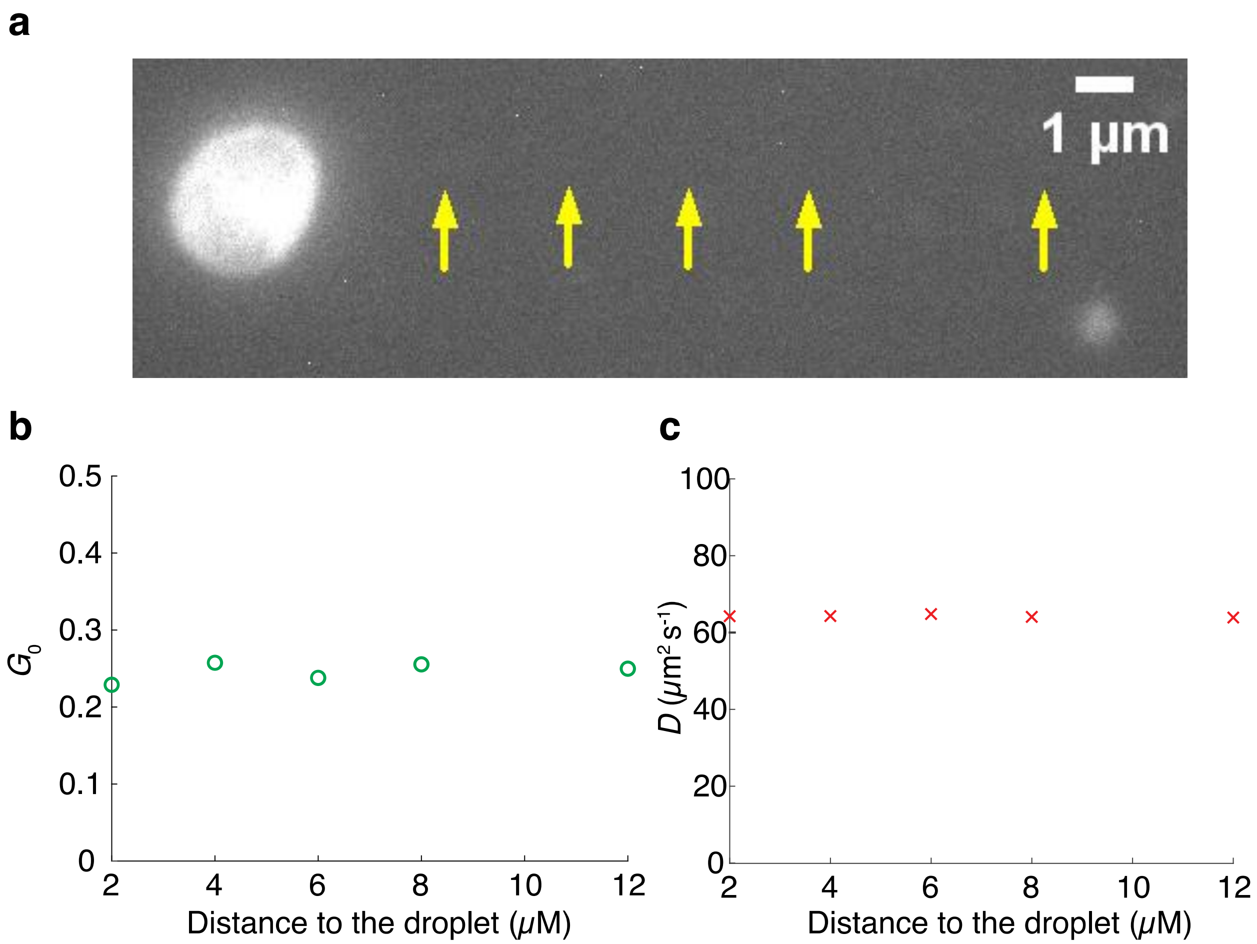


**Supplementary Figure 4 | The measured dilute-phase properties are independent of measurement location. a** FCS was performed in a focal volume 2, 4, 6, 8 and 12 $\mu$m away from the nearest droplet, as shown by yellow arrows. Bulk concentrations in the experiment: [EPYC1] = 5 $\mu$M, [Rubisco] = 0.25 $\mu$M, [EPYC1-GFP] = 40 nM. **b,c** Autocorrelation amplitude $G_0$ (**b**) and diffusion coefficient $D$ (**c**) of EPYC1-GFP fitted from the FCS data is plotted against the distance to the droplet. Both extracted parameters are highly reproducible.

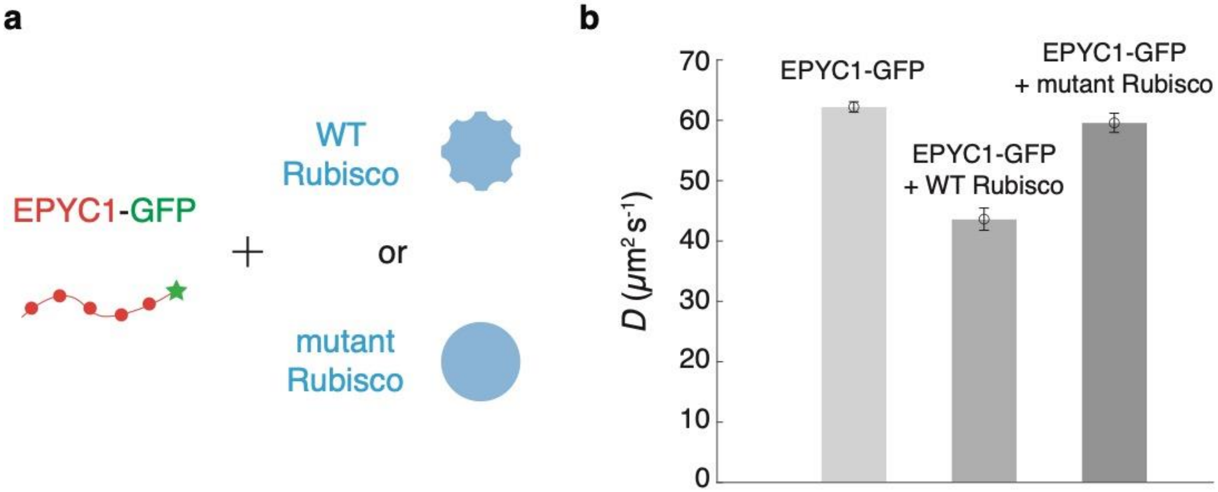


**Supplementary Figure 5 | EPYC1 does not bind to a Rubisco mutant. a** Cartoon showing the experiment setup: 10nM EPYC1-GFP is mixed with 13nM WT Rubisco or 13nM mutant Rubisco. The mutant Rubisco is M87D/V94D Rubisco small subunits from He et al. 2020[2]. **b** Diffusion rates of EPYC1-GFP alone, with 13nM WT Rubisco, or 13nM mutant Rubisco. The error bars are standard deviations of repeated experiments (*n*=5).
